# Laboratory Validation of a Clinical Metagenomic Sequencing Assay for Pathogen Detection in Cerebrospinal Fluid

**DOI:** 10.1101/330381

**Authors:** S Miller, SN Naccache, E Samayoa, K Messacar, S Arevalo, S Federman, D Stryke, E Pham, B Fung, WJ Bolosky, D Ingebrigtsen, W Lorizio, SM Paff, JA Leake, R Pesano, RL DeBiasi, SR Dominguez, CY Chiu

**Author notes:** Correspondence to:Charles Chiu Department of Laboratory Medicine and Medicine, Division of Infectious Diseases University of California, San Francisco, San Francisco, CA. These authors contributed equally to the manuscript (S Miller, SN Naccache).

## Abstract

Metagenomic next-generation sequencing (mNGS) for pan-pathogen detection has been successfully tested in proof-of-concept case studies in patients with acute illness of unknown etiology, but to date has been largely confined to research settings. Here we developed and validated an mNGS assay for diagnosis of infectious causes of meningitis and encephalitis from cerebrospinal fluid (CSF) in a licensed clinical laboratory. A clinical bioinformatics pipeline, SURPI+, was developed to rapidly analyze mNGS data, automatically report detected pathogens, and provide a graphical user interface for evaluating and interpreting results. We established quality metrics, threshold values, and limits of detection of between 0.16 – 313 genomic copies or colony forming units per milliliter for each representative organism type. Gross hemolysis and excess host nucleic acid reduced assay sensitivity; however, a spiked phage used as an internal control was a reliable indicator of sensitivity loss. Diagnostic test accuracy was evaluated by blinded mNGS testing of 95 patient samples, revealing 73% sensitivity and 99% specificity compared to original clinical test results, with 81% positive percent agreement and 99% negative percent agreement after discrepancy analysis. Subsequent mNGS challenge testing of 20 positive CSF samples prospectively collected from a cohort of pediatric patients hospitalized with meningitis, myelitis, and/or encephalitis showed 92% sensitivity and 96% specificity relative to conventional microbiological testing of CSF in identifying the causative pathogen. These results demonstrate the analytic performance of a laboratory-validated mNGS assay for pan-pathogen detection, to be used clinically for diagnosis of neurological infections from CSF.

## INTRODUCTION

Metagenomic next-generation sequencing (mNGS) provides a comprehensive method by which nearly all potential pathogens – viruses, bacteria, fungi, and parasites – can be accurately identified in a single assay (Goldberg et al. 2015; Chiu and Miller 2016). This approach is attractive for diagnosis of infectious diseases, as pathogens that cause an infectious syndrome commonly have non-specific, overlapping clinical presentations (Washington 1996). Recent advances in sequencing technology and the development of rapid bioinformatics pipelines have enabled mNGS testing to be performed within a clinically actionable time frame (Cazanave et al. 2013; Naccache et al. 2014; Wilson et al. 2014; Fremond et al. 2015; Greninger et al. 2015; Naccache et al. 2015; Salzberg et al. 2016; Mongkolrattanothai et al. 2017; Parize et al. 2017; Schlaberg et al. 2017b). However, numerous challenges remain with migrating mNGS testing into the clinical microbiology laboratory. These include (1) lack of an established blueprint for mNGS clinical validation, (2) difficulty in discriminating pathogens from colonizing microorganisms or contaminants, (3) paucity of bioinformatics software customized for clinical diagnostic use, (4) concern over quality and comprehensiveness of available reference databases, and (5) requirement for regulatory compliance inherent to patient diagnostic testing in a CLIA (Clinical Laboratory Improvement Amendments) environment.

Acute neurological illnesses such as meningitis and encephalitis are devastating syndromes, remaining undiagnosed in a majority of cases (Glaser et al. 2003; Glaser et al. 2006; Granerod et al. 2010). The diagnostic workup for many patients requires extensive, and often negative, serial testing that typically utilizes a combination of culture, antigen, serologic, and molecular methods, resulting in delayed or missed diagnoses and increased costs. Given the high burden of encephalitis-associated hospitalizations in the United States (Khetsuriani et al. 2002), there is a large unmet clinical need for better and more timely diagnostics for this syndrome, both to identify and to exclude infectious etiologies.

Here we present the development and validation of an mNGS assay for comprehensive diagnosis of infectious causes of meningitis and encephalitis from CSF, expanding on summary data presented in a previously published review (Schlaberg et al. 2017a). The analytic performance of the mNGS assay was compared to results from conventional clinical microbiological testing performed in hospital or commercial diagnostic laboratories. We also tested the assay by blinded analysis of a challenge set of 20 CSF samples prospectively collected from patients with diagnosed neurological infections at a single pediatric tertiary care hospital.

## METHODS

### mNGS Assay

The processing and analysis workflow for the mNGS assay was performed as follows (**Figure 1**), with a more detailed description provided in the **Supplemental Methods**. Briefly, each CSF sample was first subjected to bead-beating to lyse organisms (**Figure 1A)**, followed by addition (“spiking”) of DNA T1 and RNA M2 bacteriophages as an internal control (IC). Total nucleic acid was then extracted and split into 2 aliquots for construction of separate DNA and RNA libraries. Microbial sequences were enriched by antibody-based removal of methylated host DNA (for DNA libraries) or DNase treatment (for RNA libraries), followed by transposon-based library construction (**Figure 1B)**. Each sequencing run on an Illumina HiSeq instrument included up to 8 samples, along with a negative “no template” control (NTC) consisting of elution buffer, intended to allow for sensitive detection of contamination, and a positive control (PC) consisting of a mixture of 7 representative organisms (RNA virus, DNA virus, Gram-positive bacterium, Gram-negative bacterium, fungus, mold, and parasite). Some of the early sequencing data used for validation was generated on an Illumina MiSeq instrument. Sequence analysis (**Figure 1C)** using the SURPI+ computational pipeline (Naccache et al. 2014) consisted of (1) pre-processing for trimming of adapters and removal of low-complexity and low-quality reads, (2) human host background subtraction, (3) alignment to the National Center for Biotechnology Information (NCBI) GenBank NT (nucleotide) reference database for microbial identification, (4) taxonomic classification of aligned reads, and (5) visualization and interpretation of sequencing data. Receiver-operator curve (ROC) analyses were performed to determine optimal threshold values for organism detection based on mNGS data output. Each mNGS run was analyzed by experienced laboratory physicians (SM and CYC), and results were generated for 5 categories per sample (RNA virus, DNA virus, bacteria, fungi and parasite). Run quality control (QC) metrics included a minimum of 5 million reads per library, ≥100 reads per million (RPM) for the IC T1 and MS2 phages in the DNA and RNA libraries, respectively, and positive qualitative detection of each of the 7 microorganisms in the PC using pre-designated thresholds, as described below.

**Figure 1.**
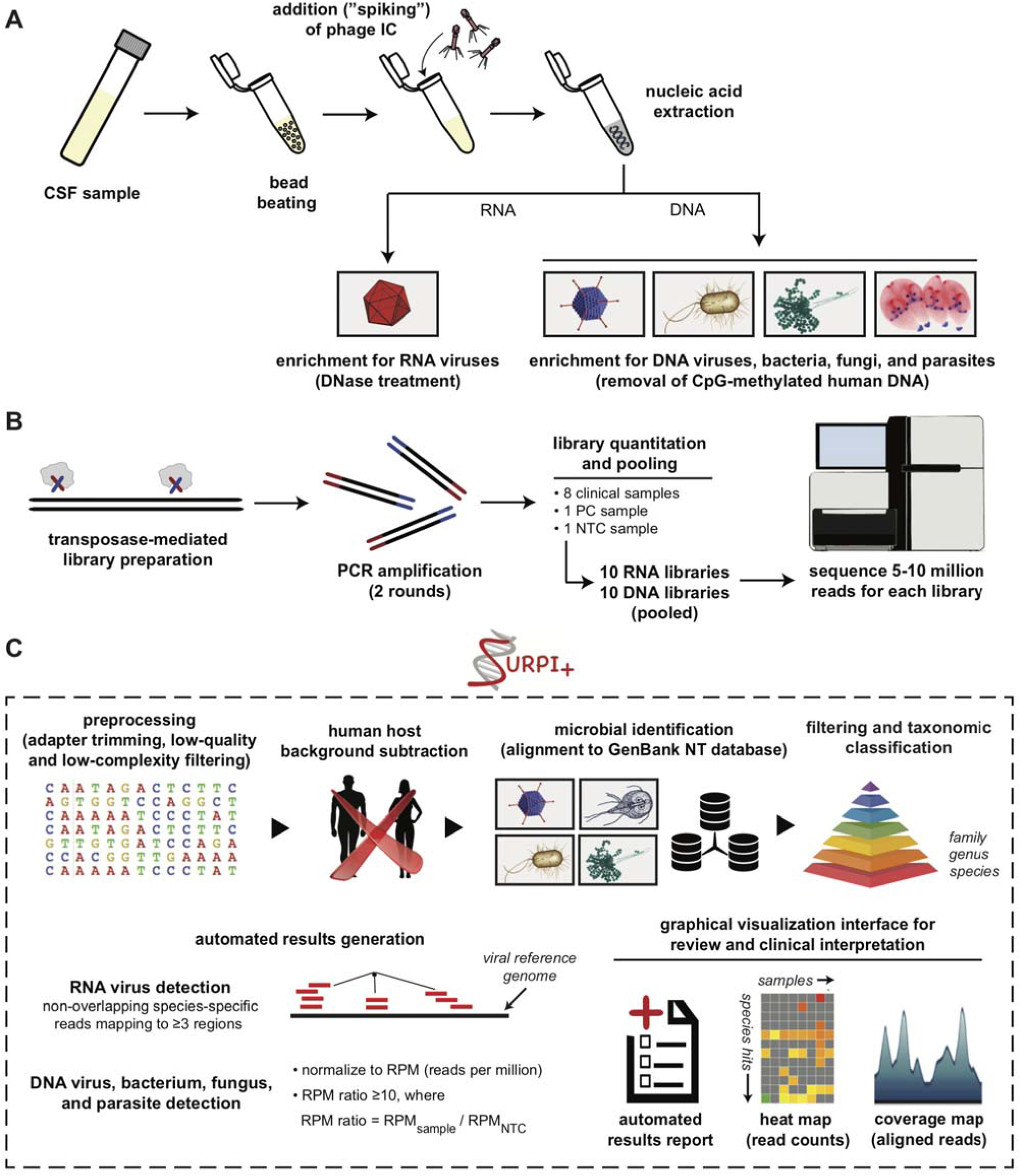
Schematic of the mNGS Assay Workflow. **(A)** CSF is extracted after lysis by bead-beating and internal control addition to allow viral, bacterial, fungal and parasite nucleic acid retrieval. Total nucleic acid extracts are enriched for pathogen DNA by removal of methylated DNA (DNA libraries) and treatment with DNase (RNA libraries). **(B)** Libraries are generated using the Nextera XT protocol and amplified using 2 rounds of PCR. Libraries are quantified, pooled, and loaded onto the sequencer. **(C)** Sequences are processed using SURPI+ software for alignment and classification. Reads are preprocessed by trimming of adapters and removal of low-quality / low-complexity sequences, followed by computational subtraction of human reads and taxonomic classification of remaining microbial reads to family, genus, or species. For viruses, reads are mapped to the closest matched genome to identify nonoverlapping regions; for bacteria, fungi, and parasites, a read per million (RPM) ratio (RPM-r) metric is calculated, defined as RPM-r = RPM_sample_ / _NTC_. To aid in analysis, result reports, heat maps of raw / normalized read counts, and coverage maps are automatically generated for use in review and clinical interpretation.

### Evaluation of mNGS Analytical Performance Characteristics

A detailed description of the methods used to evaluate mNGS analytical performance characteristics is provided in the **Supplemental Methods.** Briefly, limits of detection (LOD) were determined for each of the 7 representative organisms in the PC by analyzing a series of 10-fold dilutions for qualitative detection. The LOD was determined using probit analysis, defined as the concentration at which 95% of replicates would be detected, with at least 3 replicates performed for concentrations above and below this level. Precision was determined using repeat analysis of the PC and NTC over 20 consecutive sequencing runs (inter-assay reproducibility) and 3 sets of separately PC and NTC controls processed in parallel on the same run (intra-assay reproducibility). Test stability was determined using control samples held at various temperatures and subject to multiple freeze/thaw cycles. Interference was determined using PC spiked with known amounts of human DNA or RNA material. Results were assessed for qualitative detection of organisms in the PC.

Accuracy was determined using 95 clinical CSF samples (**Supplemental Figure S1),** with up to 5 potential result categories per sample (RNA virus, DNA virus, bacteria, fungus and parasite). Samples were obtained from patients at University of California, San Francisco (UCSF) (n=59), Children’s National Medical Center (n=19), Children’s Hospital Colorado (CHCO) (n=1), and Quest Diagnostics (n=16). Due to the varying number of clinical tests performed per sample and limited residual CSF volume, we generated 3 composite reference standards for purposes of comparison. The first composite standard consisted of the original conventional clinical test results (both positive and negative) available prior to mNGS analysis, providing an assessment of mNGS sensitivity and specificity relative to clinical testing. If mNGS detected an organism that had not been tested for clinically, the result was considered “reference untested”, and that result was excluded from the comparison. Negative mNGS results corresponding to a given category were also excluded if no clinical testing for pathogens within that category had been performed. A second composite standard consisted of combined results from the original clinical testing and additional molecular testing of CSF samples (volume permitting), either when clinical and mNGS results were discrepant (n=8) or when mNGS detected an organism that had not been included in the original testing (n=10). Finally, a third composite standard was generated, which excluded samples with high human sequence background (n=26), defined as samples with phage IC sequence recovery below a pre-designated 100 RPM threshold. The second and third comparisons are described as positive percent agreement (PPA) and negative percent agreement (NPA), as selective discrepancy testing can bias estimates of test sensitivity and specificity (Meiser 2002). To evaluate mNGS detection performance for additional organism types not readily available from clinical CSF samples, the accuracy study also included contrived samples of 5 known organisms (*N. meningitidis, S. agalactiae, C. albicans, M. fortuitum, M. abscessus*) spiked into negative CSF at defined concentrations.

### Challenge study

The Aseptic Meningitis and Encephalitis Study (AMES) is a prospective cohort study enrolling children presenting to CHCO with culture-negative meningitis and encephalitis since 2012. Ethical approval for the study was obtained from the Colorado Multiple Institutional Review Board (protocol #12-0745), and all subjects provided informed consent for specimen collection and testing. A subset of CSF specimens (n=20) with sufficient residual volume (600 uL) from subjects with known and unknown etiologies was coded for mNGS testing as a challenge set. Samples were processed in a blinded fashion at UCSF, and results discussed in clinical context with site investigators at CHCO over web-based teleconferencing. Results from the mNGS assay and conventional clinical testing were compared for up to 5 result categories (RNA virus, DNA virus, bacterium, fungus, and parasite) per sample.

### Accession numbers

Microbial sequences with human reads removed have been deposited in the NCBI Sequence Read Archive (BioProject accession number PRJNA234047). Sequences corresponding to the HIV-1 and CMV controls in the PC and the MS2 (RNA) phage and T1 (DNA) phage spiked IC samples have been deposited in GenBank (accession numbers pending).

## RESULTS

### Sample processing and bioinformatics analysis

We developed an mNGS assay for pathogen identification from CSF consisting of library preparation, sequencing, and bioinformatics analysis for pathogen detection (**Figure 1**), and validated the performance of the assay in a CLIA-certified laboratory. Wet bench protocols and sequencing runs for the study were performed by state-licensed clinical laboratory scientists. For each sequencing run, NTC (“no template” control), PC (positive control), and up to 8 patient CSF samples were processed by DNA/RNA enrichment, nucleic acid extraction, construction of DNA and RNA libraries, and sequencing on an Illumina HiSeq instrument in rapid run mode, targeting a total of 5 to 20 million sequences per library (**Figure 1A and B)**. Raw mNGS sequence data were analyzed using SURPI+, a bioinformatics analysis pipeline for pathogen identification from mNGS sequence data (Naccache et al. 2014) that was modified for clinical use. Specifically, the SURPI+ pipeline included filtering algorithms for exclusion of false-positive hits from database misannotations, taxonomic classification for accurate species-level identification, automated report generation implementing *a priori* established thresholds for pathogen detection, and a web-based graphical user interface to facilitate laboratory director review and confirmation of mNGS findings (**Figures 1C and Supplemental Figure S2).**

### Establishing thresholds for reporting detected pathogens

To minimize the potential for false positive results from low-level background contamination, threshold criteria were established for organism detection (**Figure 1C)**. For viruses, we developed threshold criteria based on the detection of nonoverlapping reads from at least 3 distinct genomic regions, taking into consideration viruses incidentally detected in the NTC sample that were potential background contaminants. Viruses comprising known flora, such as anelloviruses and papillomaviruses, or laboratory reagent contaminants, such as murine gammaretroviruses, were not reported. For identification of bacteria, fungi, and parasites, we developed a reads per million (RPM) ratio metric, or RPM-r, defined as RPM-r = RPM_sample_ / RPM_NTC_ (with the minimum RPM_NTC_ set to 1). This metric accounted for background contamination by normalizing the RPM of detected pathogen reads assigned to a given taxonomic classification (family, genus, or species) with respect to the RPM in the NTC. To determine the optimal threshold value for RPM-r, we plotted receiver operating characteristic (ROC) curves at varying ratios corresponding to mNGS analysis of 95 clinical CSF samples that were included in the accuracy evaluation (**Supplemental Figure S3**; also see below). The ROC curve analysis showed that an RPM-r of 10 maximized accuracy for organism detection. Thus, we designated a minimum threshold of 10 RPM-r for reporting the detection of a bacterium, fungus, or parasite. Occasionally, multiple bacterial genera (≥2) from environmental and/or skin flora were detected in a CSF sample above the 10 RPM-r threshold, and attributed to contamination. In these cases (n=7), mNGS results were reported as “multiple bacterial genera detected” (with an interpretive comment indicating likely sample contamination), and were considered as negative for bacterial detection by mNGS.

### Limits of detection

To calculate the 95% limits of detection (LOD), defined as the lowest concentration at which 95% of positive samples are detected, multiple replicates of the PC at serial dilutions near the estimated detection limit were tested by mNGS. Using probit analysis, a 95% limit of detection was determined for each of the 7 representative organisms in the PC (**Table 1**). The final working PC consisted of the 7 organisms spiked at concentrations ranging from 0.5-2 log above the 95% limit of detection. A linear correlation was observed between the input concentration and number of reads / genome coverage for viruses (R^2^ values 0.9083-0.9911), and between the input concentration and RPM-r for bacteria, fungi, and parasites (R^2^ values.09856-.09996) (**Supplemental Figure S4**).

**Table 1.**
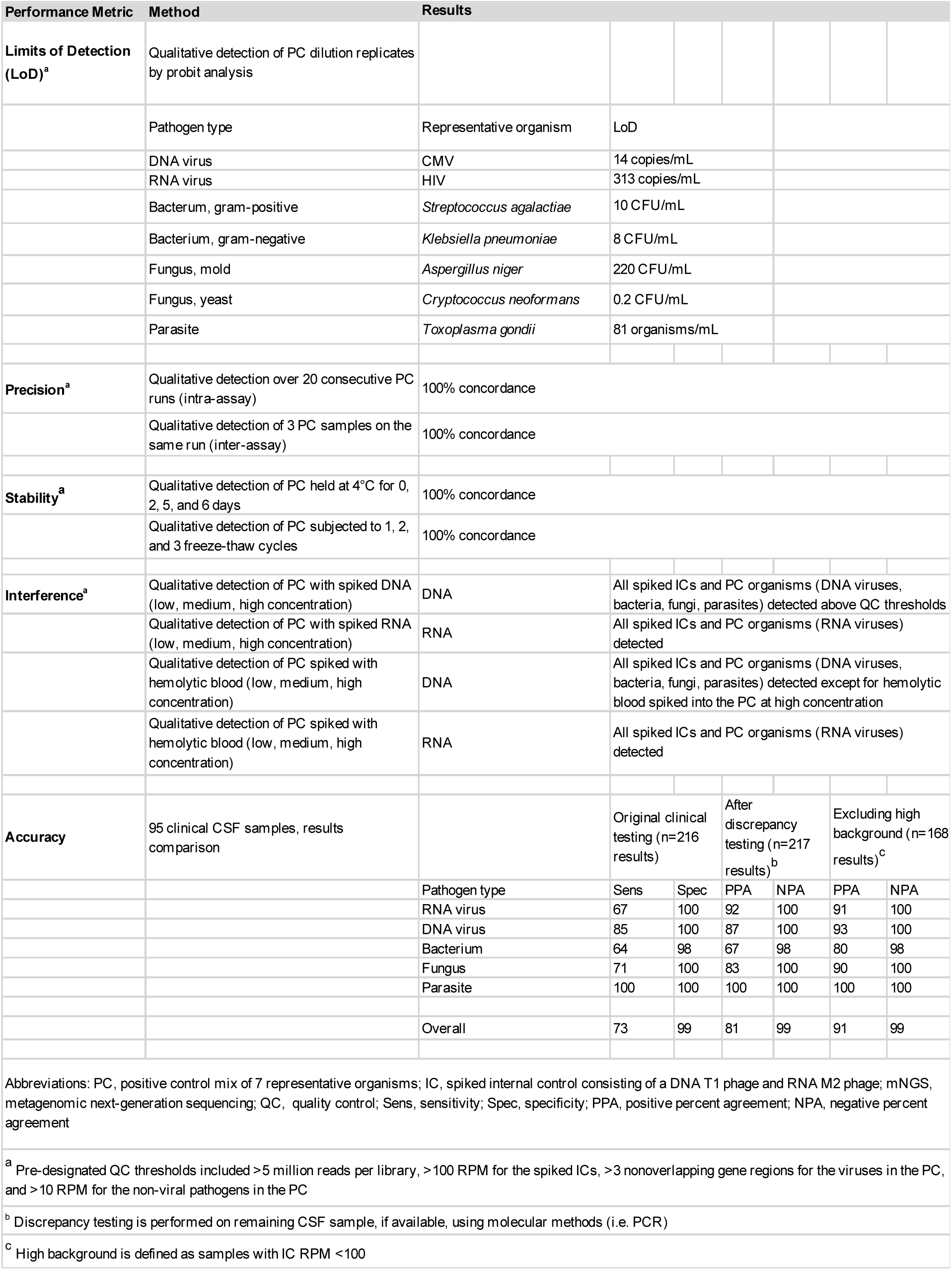
Performance characteristics for the mNGS assay

| Performance Metric | Method | Results |  |  |  |  |  |  |
| --- | --- | --- | --- | --- | --- | --- | --- | --- |
| **Limits of Detection (LoD)<sup>a</sup>** | Qualitative detection of PC dilution replicates by probit analysis |  |  |  |  |  |  |  |
|  | Pathogen type | Representative organism | LoD |  |  |  |  |  |
|  | DNA virus | CMV | 14 copies/mL |  |  |  |  |  |
|  | RNA virus | HIV | 313 copies/mL |  |  |  |  |  |
|  | Bacterium, gram-positive | *Streptococcus agalactiae* | 10 CFU/mL |  |  |  |  |  |
|  | Bacterium, gram-negative | *Klebsiella pneumoniae* | 8 CFU/mL |  |  |  |  |  |
|  | Fungus, mold | *Aspergillus niger* | 220 CFU/mL |  |  |  |  |  |
|  | Fungus, yeast | *Cryptococcus neoformans* | 0.2 CFU/mL |  |  |  |  |  |
|  | Parasite | *Toxoplasma gondii* | 81 organisms/mL |  |  |  |  |  |
| **Precision<sup>a</sup>** | Qualitative detection over 20 consecutive PC runs (intra-assay) | 100% concordance |  |  |  |  |  |  |
|  | Qualitative detection of 3 PC samples on the same run (inter-assay) | 100% concordance |  |  |  |  |  |  |
| **Stability<sup>a</sup>** | Qualitative detection of PC held at 4°C for 0, 2, 5, and 6 days | 100% concordance |  |  |  |  |  |  |
|  | Qualitative detection of PC subjected to 1, 2, and 3 freeze-thaw cycles | 100% concordance |  |  |  |  |  |  |
| **Interference<sup>a</sup>** | Qualitative detection of PC with spiked DNA (low, medium, high concentration) | DNA | All spiked ICs and PC organisms (DNA viruses, bacteria, fungi, parasites) detected above QC thresholds |  |  |  |  |  |
|  | Qualitative detection of PC with spiked RNA (low, medium, high concentration) | RNA | All spiked ICs and PC organisms (RNA viruses) detected |  |  |  |  |  |
|  | Qualitative detection of PC spiked with hemolytic blood (low, medium, high concentration) | DNA | All spiked ICs and PC organisms (DNA viruses, bacteria, fungi, parasites) detected except for hemolytic blood spiked into the PC at high concentration |  |  |  |  |  |
|  | Qualitative detection of PC spiked with hemolytic blood (low, medium, high concentration) | RNA | All spiked ICs and PC organisms (RNA viruses) detected |  |  |  |  |  |
| **Accuracy** | 95 clinical CSF samples, results comparison |  | Original clinical testing (n=216 results) |  | After discrepancy testing (n=217 results)<sup>b</sup> |  | Excluding high background (n=168 results)<sup>c</sup> |  |
|  |  | Pathogen type | Sens | Spec | PPA | NPA | PPA | NPA |
|  |  | RNA virus | 67 | 100 | 92 | 100 | 91 | 100 |
|  |  | DNA virus | 85 | 100 | 87 | 100 | 93 | 100 |
|  |  | Bacterium | 64 | 98 | 67 | 98 | 80 | 98 |
|  |  | Fungus | 71 | 100 | 83 | 100 | 90 | 100 |
|  |  | Parasite | 100 | 100 | 100 | 100 | 100 | 100 |
|  |  | Overall | 73 | 99 | 81 | 99 | 91 | 99 |

Abbreviations: PC, positive control mix of 7 representative organisms; IC, spiked internal control consisting of a DNA T1 phage and RNA M2 phage; mNGS, metagenomic next-generation sequencing; QC, quality control; Sens, sensitivity; Spec, specificity; PPA, positive percent agreement; NPA, negative percent agreement
<sup>a</sup> Pre-designated QC thresholds included >5 million reads per library, >100 RPM for the spiked ICs, >3 nonoverlapping gene regions for the viruses in the PC, and >10 RPM for the non-viral pathogens in the PC
<sup>b</sup> Discrepancy testing is performed on remaining CSF sample, if available, using molecular methods (i.e. PCR)
<sup>c</sup> High background is defined as samples with IC RPM <100

### Precision

We demonstrated inter-assay reproducibility by mNGS testing of the NTC and PC across 20 consecutive sequencing runs, and intra-assay reproducibility by testing of 3 independently generated sets of NTC and PC on the same run. Internal spiked phage controls passed QC for every run, and only one PC RNA library (out of 46 total DNA and RNA libraries) had fewer than the minimum designated cutoff of 5 million reads. All 7 organisms were detected using pre-established threshold criteria for the intra-assay run and each replicate inter-assay run (**Table 1**).

### Accuracy

For evaluation of accuracy, a total of 95 CSF patient samples (73 positive and 22 negative for a pathogen by conventional clinical testing) were tested using the mNGS assay. There were 5 categories of results for each sample, corresponding to 5 different pathogen types (bacteria, fungi, parasites, DNA viruses, and RNA viruses). mNGS results were compared to (1) original clinical test results, (2) results after discrepancy testing, and (3) results after discrepancy testing and exclusion of samples with high host background (**Table 1 and Supplemental Table S1**).

Overall, the mNGS assay showed 73% sensitivity and 99% specificity compared to original clinical test results. 21 mNGS results were considered false-negatives, including 4 RNA viruses (1 enterovirus and 3 West Nile virus (WNV) diagnosed by CSF serology), 4 DNA viruses (2 VZV, 1 HSV-2, 1 EBV, diagnosed by PCR), 9 bacteria (diagnosed by culture), and 4 fungi (diagnosed by culture and/or antigen testing) (**Supplemental Table S1**). One mNGS result was considered to be false-positive for *Bacillus* sp. detection in a culture-negative CSF. All 5 organisms spiked into negative CSF matrix as part of the accuracy study were correctly detected (*N. meningitidis, S. agalactiae, C. albicans, M. fortuitum, M. abscessus*).

Discrepancy analysis using targeted clinical PCR was performed on 18 samples with sufficient volume available for testing. For organisms detected by mNGS but not tested for clinically (n=10), discrepancy tests confirmed the mNGS results in all 10 cases (5 HIV, 1 CMV, 2 EBV, 1 HSV, 1 HHV6). In 8 cases, mNGS results were initially considered to be false negatives but enough sample was available for discrepancy testing using molecular methods. Overall, discrepancy testing using molecular testing failed to detect the causative organism in 5 of the 8 cases with negative mNGS results.Two cases of WNV diagnosed by positive CSF IgM serology (1 case also had cross-reactive IgM antibodies to Japanese Encephalits Virus (JEV) in blood) were negative by WNV PCR testing, concordant with the mNGS results. Two culture-positive bacterial cases with negative mNGS results underwent orthogonal testing using 16S rRNA bacterial PCR, and one was not detected (*P. mirabilis*), whereas one was positive (*E. galinarum*). Four culture-positive samples with negative mNGS results for fungi were tested using 18S internal transcribed spacer (ITS) PCR, and 2 were negative (*C. parapsilosis* and *C. neoformans*) whereas the other 2 were positive (*A. fumigatus* and *S. schenkii*).

Among the 3 remaining *bona fide* false-negative mNGS cases (*E. galinarum, A. fumigatus,* and *S. schenkii*), the first 2 cases were only weakly positive by original clinical testing (the *E. galinarum* grew from broth only, whereas the *A. fumigatus* was galactomannan-positive and fungal culture negative). Both were also high-background samples **(Supplemental Table S1),** and thus the number of identified pathogen reads did not meet thresholds for reporting. The third case (*S. schenckii*) was likely missed by mNGS testing because the ∼32 Mb genome of *S. schenckii*, while publicly available (Cuomo et al. 2014), is not part of the GenBank NT reference database used by the SURPI+ computational pipeline. This resulted in only 33 reads in the sample (RPM-r 1.93) being identified as *S. schenckii*, also below reporting thresholds. Adjusting the results comparison on the basis of the discrepancy testing results yielded 81% positive percent agreement and 99% negative percent agreement for the mNGS assay relative to combined original and discrepancy testing results.

There were additional incidental organism detections by mNGS (n=12) where sample volume was insufficient to perform confirmatory molecular testing. These included HIV (n=4), WNV (n=1), rotavirus (n=1), rhinovirus (n=1), HCV (n=1), parvovirus B19 (n=2), human herpesvirus 7 (n=1) and *Bacillus* spp. (n=1). With the exception of *Bacillus* spp. which might have been recovered in CSF cultures, the presence of these organisms could not be independently confirmed by molecular testing and thus these mNGS results were excluded from the comparisons, as it could not be determined whether a given additional detection was a true or false positive.

A third comparison was performed after exclusion of results from CSF samples with an IC RPM of <100, indicating potential decreased sensitivity for the mNGS assay due to high human host background (see “Interference”, below). A total of 26 samples had high background (1 RNA virus, 3 DNA virus, 19 bacteria, 2 fungi, 1 negative), and exclusion of these yielded 91% positive percent agreement and 99% negative percent agreement for the mNGS assay overall. Notably, the 19 bacterial samples with high background comprised 70.4% of the total number of culture-positive bacterial cases (n=27), consistent with the relatively high leukocyte levels seen in typical cases of bacterial meningitis.

### Interference

We evaluated the effects of interference with human DNA and RNA, red blood cell hemolysis, and mixtures of related species in the same genus (*Staphylococcus aureus* and *Staphylococcus epidermidis*) on mNGS assay performance **(Table 1)**. Addition of human DNA at a level equivalent to 1 x 10^6^ cells/mL resulted in complete failure to detect spiked DNA pathogens in the PC, whereas addition of exogenous RNA and DNA at lower levels (≤1 x 10^4^ cells/mL) did not impact qualitative detection. The number of sequenced IC phage reads was found to be linearly correlated with the amount of added exogenous DNA (R^2^ = 0.999). Based on the interference results, an RPM threshold of 100 was chosen for the IC phage reads, with RPM values below this level indicating that the sample had high human DNA and/or RNA background (**Supplemental Figure S5**). For mNGS reporting, these high-background samples included a comment that the assay had decreased sensitivity for detection of RNA viruses (from RNA libraries) or DNA viruses, bacteria, fungi and parasites (from DNA libraries).

Available data from 55 CSF samples in the accuracy study with recorded white blood cell (WBC) counts were used to evaluate the effect of WBC count, related to the amount of human nucleic acid background, on recovery of IC phage sequences. Among 26 samples with IC DNA phage counts of <100 RPM, indicating high human background, the average WBC was 5,896 cells/mm^3^, while 29 samples with IC counts of >100 RPM had an average WBC count of 27 cells/ mm^3^ (p = 0.0498 by two-tailed t-test).

Gross hemolysis (dark red CSF) resulted in decreased sensitivity for RNA virus detection (HIV-1 in the PC) by mNGS, but did not affect detection sensitivity for DNA pathogens. Moderate to low levels of hemolysis (pink to light red CSF) did not affect detect sensitivity for any of the PC organisms. Analysis of spiked samples containing *S. aureus* and *S. epidermidis* with equivalent RPM-r values at baseline demonstrated accurate discrimination of species within the same genus when mixed at 1:1, 4:1, and 1:4 ratios, as both species were correctly identified and calculated RPM-r values were within 7% of that expected on the basis of the spiked amounts.

### Stability

Analysis of replicates of the PC held at 4°C for 0, 2, 5, and 6 days and subjected to 3 freeze-thaw cycles demonstrated detection of all organisms **(Table 1)**.

### Challenge study

We blindly evaluated the performance of the mNGS assay on a set of 20 prospectively collected CSF samples from pediatric patients hospitalized at CHCO with meningitis, encephalitis, and/or myelitis (**Supplemental Table S2**). Comparison of the results assembled from each of the 5 organism categories yielded a sensitivity of 92% and specificity of 96% for mNGS testing relative to conventional microbiological testing of CSF (culture, PCR, antigen, and serological testing). The assay correctly identified the causative pathogen in 11 of 12 cases that were previously positive for direct organism detection or serology from CSF, including cases of enterovirus (n=8), HSV-1 (n=1), HIV-1 (n=1) and WNV (n=1) in a patient with positive IgM serology from CSF. The mNGS assay failed to detect WNV in a second patient with positive CSF IgM serology. Three additional organisms (*Enterobacter sp. Corynebacterium sp.,* and EBV) were detected by mNGS, each from a different sample. The detection of *Enterobacter sp.* and *Corynebacterium sp.* were considered to be mNGS false-positives, since the samples had previously tested negative by culture. mNGS also identified EBV in a CSF sample from a patient with positive testing for EBV IgG antibodies in blood; this finding was excluded from the comparison due to the lack of confirmatory testing from CSF. In addition, mNGS failed to detect organisms in 4 cases: 1 case of *Borrelia burgdorferi* diagnosed using peripheral blood serology only, 2 cases of presumptive *Mycoplasma* encephalitis with PCR-positive respiratory but not CSF samples, and 1 case of presumptive enterovirus 71 infection with a positive viral culture from rectal swab but negative CSF PCR. Since the diagnoses in these 4 cases were not made directly from CSF, these results were also excluded from the comparison. Negative mNGS testing in 4 undiagnosed cases was concordant with negative conventional microbiological testing, including 1 case of culture-negative, presumptive bacterial meningitis and 3 cases of idiopathic encephalitis.

## DISCUSSION

Here we developed and validated a clinical mNGS assay intended to diagnose infectious etiologies of meningitis, encephalitis, and myelitis from CSF, followed by blinded evaluation of mNGS performance using a set of 20 prospectively collected CSF samples from pediatric patients admitted to a tertiary care hospital. As CSF is considered a normally sterile site, we postulated that mNGS data generated from testing of this body fluid type would be more straightforward to interpret than data from more “environmental” samples such as respiratory secretions and stool. However, numerous challenges had to be overcome for successful implementation of mNGS in the clinical laboratory. First, a universal sequencing library preparation protocol was required that was robust across the wide range of potential nucleic acid concentrations in patient CSF (0 – 10^8^ cells/mm^3^). This ultimately required a protocol incorporating two PCR steps, an initial step for library amplification and a recovery amplification step to ensure robust library construction from relatively acellular CSF samples or the NTC buffer control, both containing little to no human host background. Second, the mNGS assay had to be capable of simultaneously detecting a broad spectrum of pathogens, including viruses (both single- and double-stranded RNA and DNA genomes), bacteria, fungi, and parasites.

Thus, the mNGS protocol incorporated (1) a bead-beating step for lysis of microbial cell walls, (2) separate construction of RNA and DNA libraries from nucleic acid extracts for detection of RNA viruses and DNA pathogens, respectively, and (3) bioinformatics analysis using the entirety of NCBI GenBank NT database as a comprehensive reference database. Finally, reproducible threshold metrics needed to be developed and evaluated using ROC curve analysis to enable correct identification of pathogens from mNGS data above background noise.

We developed quality control materials and metrics for the CSF mNGS assay, including acceptable criteria for the performance of external positive and negative controls, as well as spiked internal controls. Given the untargeted nature of mNGS, a key limitation of the approach for infectious disease diagnostics is background interference, generally from human host DNA. The use of a spiked phage IC was found to be useful for assessing whether high background was present, indicating decreased sensitivity of pathogen detection by mNGS (Schlaberg et al. 2017a). Overall, 27.4% of DNA libraries and 6.3% of RNA libraries in the accuracy study had fewer than 100 RPM IC phage reads recovered, making background interference a fairly common limitation. Thus, in high background samples, negative mNGS findings may be less useful for excluding infection, and other diagnostic tests that may be less sensitive to background should be considered, such as 16S rRNA bacterial PCR (Salipante et al. 2013) and ITS fungal PCR (Pryce et al. 2006). This is especially relevant in cases of bacterial meningitis with high leukocyte counts in CSF. However, despite this limitation, mNGS was still able to detect bacterial pathogens in 12 of 19 culture-positive samples in the accuracy study with high host background.

The overall accuracy of the mNGS assay for pathogen detection over 5 categories of microorganisms as compared to initial conventional microbiological testing was 90%, with 73% sensitivity and 99% specificity. Positive percent agreement rose to 81% after discrepancy testing of samples with sufficient volume, and exclusion of samples with high host background increased this further to 91%. Only a fraction of all possible diagnostic tests for pathogens are performed in clinical microbiology laboratories given limited CSF sample volume. Thus, we decided to exclude organism detections by mNGS for which independent testing had not been performed in assessment of initial test performance (n=22), as no reference result was available. However, in 10 cases with sufficient CSF volume for orthogonal confirmatory testing, all 10 were found to be analytical true positives. Furthermore, in 5 of 8 (62.5%) cases where mNGS failed to detect an organism that was found with initial clinical testing (using culture and/or serology), confirmatory orthogonal PCR testing for this organism was negative, indicating that the sample may have degraded over time or that the original clinical result may have been incorrect.

As with any diagnostic assay, mNGS testing is prone to contamination. There is the potential to detect colonizing organisms that constitute normal human body flora (e.g. anelloviruses in CSF and blood (Maggi and Bendinelli 2010; Moustafa et al. 2017)), as well as exogenous (Strong et al. 2014) or cross-contamination. Often, the identity of the species detected can provide clues as to the contamination source such as skin flora (e.g. *S. epidermidis*, papillomaviruses), laboratory reagents (murine gammaretroviruses, *E. coli*, insect viruses), body flora (e.g. anelloviruses), or environmental flora (e.g. *Thermus* sp., *Bacillus* sp.). Cross-contamination in particular is a major concern given that the mNGS protocol involves PCR amplification. Strict processing controls to minimize contamination are essential and include unidirectional workflow, positive pressure ventilation in pre-amplification areas, and workspace separation for different assay steps. New reagent lots must undergo QC testing with mNGS of a reference standard, such as a previously run sample, before reagents can be put into clinical use. Background contamination is also continually monitored by keeping track of contaminants seen in the NTC or PC, and conservative threshold criteria are used to minimize the reporting of false-positive results. Additionally, periodic swipe tests of instruments and lab sources followed by mNGS of the swabs can facilitate targeted cleaning to ensure absence of laboratory contamination. However, despite use of contamination controls for the mNGS assay, 7.4% of clinical samples in the accuracy study had multiple bacteria genera detected above thresholds, generally consisting of environmental or skin flora. Rarely are bacterial co-infections causative for cases of meningitis / encephalitis (with the possible exception of brain abscesses communicating with CSF), so these findings were noted as indicating probable sample contamination, and were considered as negative for bacterial pathogen detection in this analysis. It is likely that bacterial DNA is introduced from *bona fide* uncultivatable organisms during sample collection or via reagents / tubes that are sterile but not DNA-free, and mNGS analysis and interpretation must be able to deal with these contamination risks.

The challenge study demonstrated that mNGS detected the same organism identified through conventional direct organism detection methods or serology from CSF in 11 of 12 (91.7%) of cases. One case of WNV, diagnosed serologically, was missed by mNGS (false-negative). There were 2 mNGS false-positives (*Corynebacterium sp.* and *Enterobacter sp.*), likely due to contamination introduced during sample collection, handling, or the assay procedure. mNGS of CSF failed to identify the pathogen in 4 patients who had a presumptive infectious diagnosis from peripheral microbiological testing (serology, culture, and/or PCR) done from sites other than CSF. We found that the sensitivity of mNGS is critically dependent on whether the organism (or nucleic acid from the organism) is present at the time of sample collection. It is thus not unexpected that mNGS testing missed a number of infections that are most often diagnosed by serology because the pathogen is absent or only transiently present in CSF (e.g. neurosyphilis, Lyme neuroborreliosis, and WNV). For these cases, direct detection testing approaches such as PCR and mNGS may be inappropriate and there should be a low threshold for ordering antibody-based serologic testing or performing microbial analysis from other body sites to establish the diagnosis if there is clinical suspicion (Debiasi and Tyler 2004).

While mNGS testing can provide broad-spectrum pathogen identification, assessment of the clinical significance of the findings requires interpretation. Thus, mNGS results are submitted to the patient electronic medical record as an interpretive report by pathologists with expertise in microbiology and genomics, after reviewing and citing of relevant literature. In addition to the submitted report, direct discussion or teleconferences can also be set up between pathologists and providers to clarify and review mNGS results in clinical context. These forums can also be used to communicate results of supplementary analyses of mNGS data, including (1) genome assembly for characterization of predicted antibiotic or antiviral resistance mutations, (2) phylogenetic analysis for genotyping and strain-level identification, and (3) disclosure of reads from potential pathogens below formal reporting thresholds. Our finding of a linear correlation between the number of reads or genome coverage and pathogen titer (as previously noted for influenza virus in nasal swabs (Greninger et al. 2010)) also raises the prospect of extracting quantitative information from metagenomic sequence data. The metagenomic analyses performed here were facilitated by the use of SURPI+ software, as it provides summaries and and graphical visualization tools tailored for evaluation and reporting of mNGS results. Thus, the clinical relevance of mNGS findings can be efficiently communicated to physicians, potentially informing the next steps in diagnosis, management, and treatment of the patient, and may also prove informative for public health surveillance and outbreak investigation (Chiu et al. 2017).

## AUTHOR CONTRIBUTIONS

CYC SM ES and SNN developed the project. ES EP SA BF WL generated libraries. SNN SM SM CYC analyzed data for clinical validation. SF DS CYC developed SURPI+ software and graphical user interface for clinical use. WB modified the SNAP algorithm to facilitate taxonomic classification by SURPI+, SP banked CSF samples. BG JAL SD KM SM CYC provided clinical specimens. KM SD SM CYC JAL SNN SP CYC SM conducted chart review. DI BF SA conducted discrepancy testing. SM SNN and CYC wrote the manuscript with contributions from all authors.

## ACKNOWLEDGEMENTS

We thank the staff of the Clinical Immunology lab for their help in discrepancy testing, and Gail Cunningham for her expert help in obtaining AFB organisms. We would also like to thank Brittany Goldberg for providing clinical samples for use in the validation.

This work is supported by NIH grants R01 HL105704 (CYC), a UC Center for Accelerated Innovation grant funded by NIH grants U54 HL119893 and NCATS UCSF-CTSI grant UL1 TR000004 (CYC), the California Initiative to Advance Precision Medicine (CYC and SM), research support from Abbott Laboratories, Inc (CYC), and supplemental funding from the UCSF Medical Center (CYC and SM).

